# An open library of human kinase domain constructs for automated bacterial expression

**DOI:** 10.1101/038711

**Authors:** Steven K. Albanese, Daniel L. Parton, Mehtap Işık, Lucelenie Rodríguez-Laureano, Sonya M. Hanson, Julie M. Behr, Scott Gradia, Chris Jeans, Nicholas M. Levinson, Markus A. Seeliger, John D. Chodera

## Abstract

Kinases play a critical role in many cellular signaling pathways and are dysregulated in a number of diseases, such as cancer, diabetes, and neurodegeneration. Since the FDA approval of imatinib in 2001, therapeutics targeting kinases now account for roughly 50% of current cancer drug discovery efforts. The ability to explore human kinase biochemistry, biophysics, and structural biology in the laboratory is essential to making rapid progress in understanding kinase regulation, designing selective inhibitors, and studying the emergence of drug resistance. While insect and mammalian expression systems are frequently used for the expression of human kinases, bacterial expression systems are superior in terms of simplicity and cost-effectiveness but have historically struggled with human kinase expression. Following the discovery that phosphatase coexpression could produce high yields of Src and Abl kinase domains in bacterial expression systems, we have generated a library of 52 His-tagged human kinase domain constructs that express above 2 *µ*g/mL culture in a simple automated bacterial expression system utilizing phosphatase coexpression (YopH for Tyr kinases, Lambda for Ser/Thr kinases). Here, we report a structural bioinformatics approach to identify kinase domain constructs previously expressed in bacteria likely to express well in a simple high-throughput protocol, experiments demonstrating our simple construct selection strategy selects constructs with good expression yields in a test of 84 potential kinase domain boundaries for Abl, and yields from a high-throughput expression screen of 96 human kinase constructs. Using a fluorescence-based thermostability assay and a fluorescent ATP-competitive inhibitor, we show that the highest-expressing kinases are folded and have well-formed ATP binding sites. We also demonstrate how the resulting expressing constructs can be used for the biophysical and biochemical study of clinical mutations by engineering a panel of 48 Src mutations and 46 Abl mutations via single-primer mutagenesis and screening the resulting library for expression yields. The wild-type kinase construct library is available publicly via Addgene, and should prove to be of high utility for experiments focused on drug discovery and the emergence of drug resistance.

## Introduction

Kinases play a critical role in cellular signaling pathways, controlling a number of key biological processes that include growth and proliferation. There are over 500 kinases in the human genome^1, 2^, many of which are of therapeutic interest. Perturbations due to mutation, translocation, or upregulation can cause one or more kinases to become dysregulated, often with disastrous consequences^3^. Kinase dysregulation has been linked to a number of diseases, such as cancer, diabetes, and inflammation. Cancer alone is the second leading cause of death in the United States, accounting for nearly 25% of all deaths; in 2015, over^1^.7 million new cases were diagnosed, with over 580,000 deaths^4^. Nearly 50% of cancer drug development is targeted at kinases, accounting for perhaps 30% of *all* drug development effort globally^5, 6^.

The discovery of imatinib, an inhibitor that targets the Abelson tyrosine kinase (Abl) dysregulated in chronic myelogenous leukemia (CML) patients, was transformative in revealing the enormous therapeutic potential of selective kinase inhibitors, kindling hope that this remarkable success could be recapitulated for other cancers and diseases^7^. While there are now 39 FDA-approved selective kinase small molecule inhibitors (as of 16 Jan 2018)^8, 9^, these molecules were approved for targeting only 22 out of ~500 human kinases^1^, with the vast majority developed to target just a handful of kinases^10^. The discovery of therapeutically effective inhibitors for other kinases has proven remarkably challenging.

While these inhibitors have found success in the clinic, many patients cease to respond to treatment due to resistance caused by mutations in the targeted kinase^11^, activation of downstream kinases 3, or relief of feedback inhibition in signaling pathways^12^. These challenges have spurred the development of a new generation of inhibitors aimed at overcoming resistance^13, 14^, as well as mutant-specific inhibitors that target kinases bearing a missense mutation that confers resistance to an earlier generation inhibitor^15^. The ability to easily engineer and express mutant kinase domains of interest would be of enormous aid in the development of mutant-selective inhibitors, offering an advantage over current high-throughput assays^16–18^, which typically include few clinically-observed mutant kinases.

Probing human kinase biochemistry, biophysics, and structural biology in the laboratory is essential to making rapid progress in understanding kinase regulation, developing selective inhibitors, and studying the biophysical driving forces underlying mutational mechanisms of drug resistance. While human kinase expression in baculovirus-infected insect cells can achieve high success rates^19, 20^, it cannot compete in cost, convenience, or speed with bacterial expression. *E. coli* expression enables production of kinases without unwanted post-translational modifications, allowing for greater control of the system. A survey of 62 full-length non-receptor human kinases found that over 50% express well in *E. coli*^19^, but often expressing only the soluble kinase domains are sufficient, since these are the molecular targets of therapy for targeted kinase inhibitors and could be studied even for receptor-type kinases. While removal of regulatory domains can negatively impact expression and solubility, coexpression with phosphatase was shown to greatly enhance bacterial kinase expression in Src and Abl tyrosine kinases, presumably by ensuring that kinases remain in an unphosphorylated inactive form where they can cause minimal damage to cellular machinery^21^.

The protein databank (PDB) now contains over 100 human kinases that were expressed in bacteria, according to PDB header data. Many of these kinases were expressed and crystallized as part of the highly successful Structural Genomics Consortium (SGC) effort to increase structural coverage of the human kinome^22^. Since bacterial expression is often complicated by the need to tailor construct boundaries, solubility-promoting tags, and expression and purification protocols individually for each protein expressed, we wondered whether a simple, uniform, automatable expression and purification protocol could be used to identify tractable kinases, select construct boundaries, express a large number of human kinases and their mutant forms, and produce a convenient bacterial expression library to facilitate kinase research and selective inhibitor development. As a first step toward this goal, we developed a structural informatics pipeline to use available kinase structural data and associated metadata to select constructs from available human kinase libraries to clone into a standard set of vectors intended for phosphatase coexpression under a simple automatable expression and purification protocol. Using an expression screen for multiple construct domain boundaries of Abl, we found that transferring construct boundaries from available structural data can produce constructs with useful expression levels, enabling simple identification of construct domain boundaries. We then completed an automated expression screen in Rosetta2 cells of 96 different kinases and found that 52 human kinase domains express with yields greater than 2 *µ*g/mL culture. To investigate whether these kinases are properly folded and useful for biophysical experiments, we performed a fluorescence-based thermostability assay on the 14 highest expressing kinases in our panel and a single-well high-throughput fluorescence-based binding affinity measurement on 39 kinases. These experiments demonstrated that omany of the expressed kinases were folded, with well formed ATP binding sites capable of binding a small molecule kinase inhibitor. To demonstrate the utility of these constructs for probing the effect of clinical mutations on kinase structure and ligand binding, we subsequently screened 48 Src and 46 Abl mutations, finding that many clinically-derived mutant kinase domains can be expressed with useful yields in this uniform automated expression and purification protocol.

All source code, data, and wild-type kinase plasmids associated with this project are freely available online:

- **Source code and data**: https://github.com/choderalab/kinase-ecoli-expression-panel
- **Interactive table of expression data**: http://choderalab.org/kinome-expression
- **Plasmids**: https://www.addgene.org/kits/chodera-kinase-domains

## Results

### Construct boundary choice impacts Abl kinase domain expression

To understand how alternative choices of expression construct boundaries can modulate bacterial expression of a human kinase domain, we carried out an expression screen of 84 unique construct boundaries encompassing the kinase domain of the tyrosine protein kinase ABL1.

Three constructs known to express in bacteria were chosen from the literature and used as controls, spanning Uniprot residues 229-500 (PDBID: 3CS9)^23^, 229-512 (PDBID: 2G2H)^24^ and 229-515 (PDBID: 2E2B)^25^. 81 constructs were generated combinatorially by selecting nine different N-terminal boundaries spanning residues 228-243 and nine different C-terminal boundaries spanning residues 490-515, chosen to be near the start and end points for the control constructs (Figure 1A). Each of the three control constructs included six replicates to provide an estimate of the typical standard error in expression readout for the experimental constructs, which was found to be between 0.42-1.5 *µ*g/mL (Figure 1A, green constructs).

**Figure 1.**
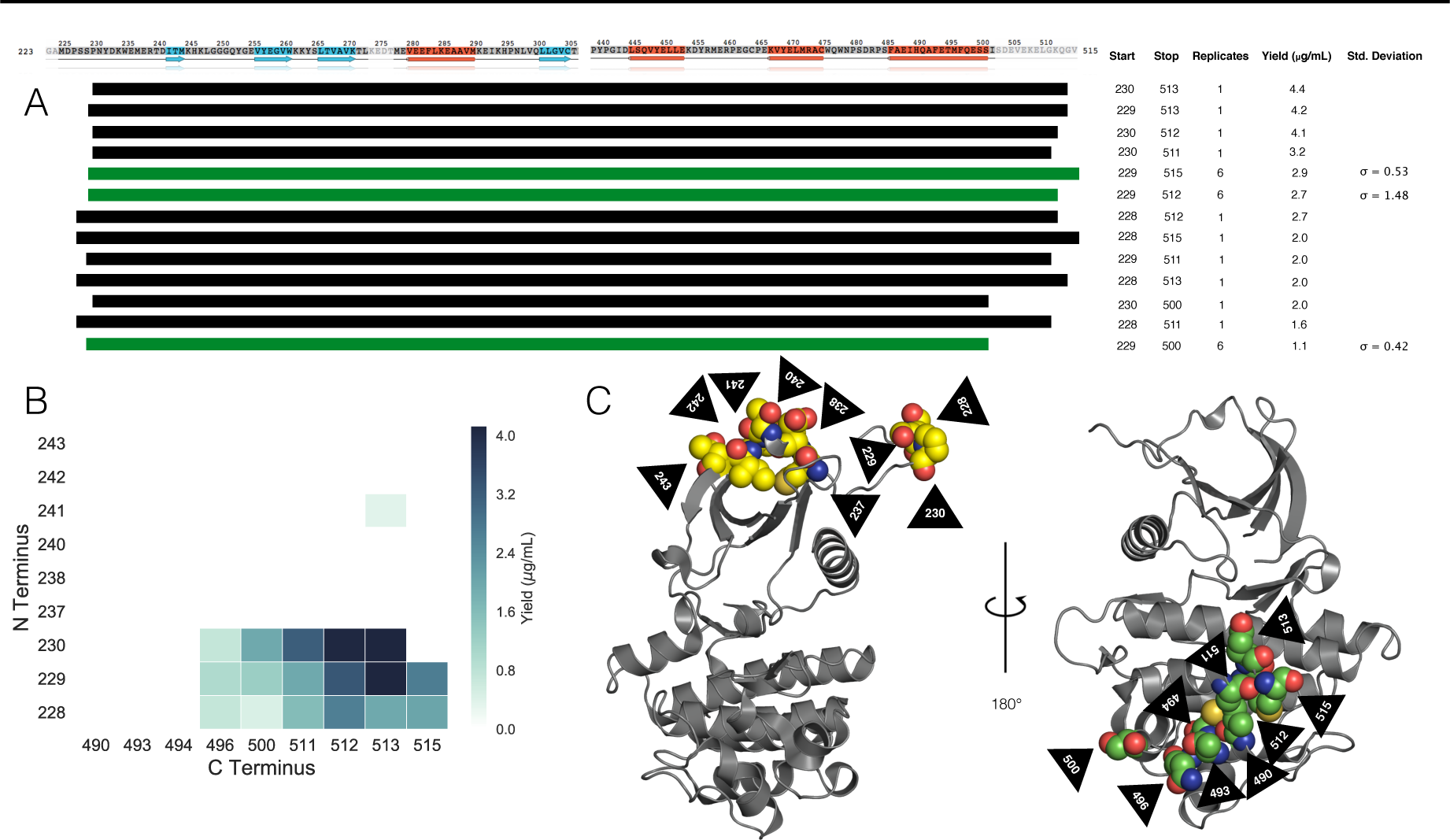
Abl kinase domain construct expression screen illustrates high sensitivity to construct boundaries. (**A**) Abl kinase domain construct boundaries with highest expression yields. Standard deviations of the yield are listed for control constructs for which six replicates were performed to give an indication of the uncertainty in experimental constructs. Secondary structure is indicated on the sequence. Beta sheets are colored blue and alpha helices are colored orange. (**B**) Heatmap showing average yields for constructs (in *µ*g/mL culture) with detectable expression as a function of N- and C-terminal construct boundaries. (**C**) *left*: PDBID: 2E2B with the nine N-terminal construct boundary amino acids shown as yellow spheres. *right*: PDBID: 4XEY with the nine C-terminal construct boundary amino acids shown as green spheres. Black arrows indicate residue numbers.

Briefly, the impact of construct boundary choice on Abl kinase domain expression was tested as follows (see Methods for full details). His10-TEV N-terminally tagged wild-type Abl constructs^2^ were coexpressed with YopH phosphatase in a 96-well format with control replicates distributed randomly throughout the plate. His-tagged protein constructs were recovered via a single nickel affinity chromatography step, and construct yields were quantified using microfluidic capillary electrophoresis following thermal denaturation. Expression yields are summarized in Figure 1A, and a synthetic gel image from the constructs with detectable expression is shown in Figure 2. Abl construct bands are present at sizes between 29 and 35 kDa (due to the variation in construct boundaries), and YopH phosphatase (which is not His-tagged but has substantial affinity for the nickel beads) is present in all samples at its expected size of 50 kDa. Strikingly, despite the fact that N-terminal and C-terminal construct boundaries only varied over 15-25 residues, only a small number of constructs produced detectable expression (Figure 1B). As highlighted in Figure 1C (left), the best N-terminal boundaries (residues 228, 229, 230) are located on a disordered strand distant from any secondary structure; N-terminal boundaries closer to the beta sheet of the N-lobe gave poor or no detectable expression (Figure 1B).

**Figure 2.**
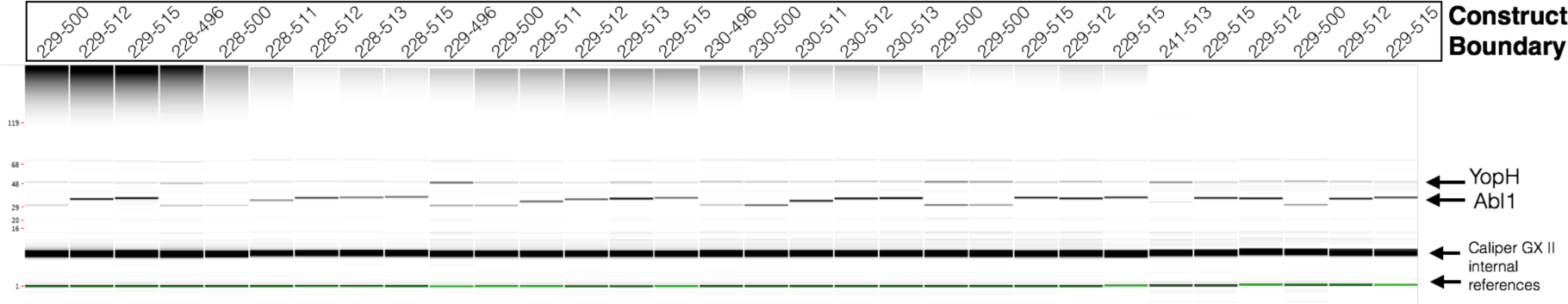
Expression yields of Abl kinase domain constructs for all constructs with detectable expression. A synthetic gel image rendering generated from Caliper GX II microfluidic gel electrophoresis data following Ni-affinity purification and thermal denaturation for all Abl constructs with detectable expression. Each well is marked with the Abl kinase domain construct residue boundaries (Uniprot canonical isoform numbering). Bands for YopH164 phosphatase (50 kDA) and Abl kinsase domain constructs (28-35 kDA) are labeled.

The best C-terminal construct boundaries (residues 511 and 512) occur in an *α*-helix (Figure 1C, right). Of note, this *α*-helix is not resolved in PDBID:2E2B^25^, suggesting this structural element may only be weakly thermodynamically stable in the absence of additional domains. In previous work, this *α*-helix was shown to undergo a dramatic conformational change which introduces a kink at residue 516, splitting the *α*-helix into two^26^. This suggests a high potential for flexibility in this region.

Two of the control constructs (which differ in construct boundary by only one or two residues) were in the top six expressing constructs (Figure 1A), and were in fact within 60% of the maximum observed expression yield. From this, we concluded that transferring construct boundaries from existing kinase domain structural data would be sufficient to bias our constructs toward useful expression levels for a large-scale screen of multiple kinases.

### Screen of 96 kinases finds 52 with useful levels of automated *E. coli* expression

To begin exploring which human kinase domains can achieve useful expression in *E. coli* using a simple automatable expression and purification protocol, a panel of kinase domain constructs for 96 kinases, for which bacterial expression has been previously demonstrated, was assembled using a semi-automated bioinformatics pipeline. Briefly, a database was built by querying Uniprot^27^ for human protein kinase domains that were both active and not truncated. This query returned a set of target sequences that were then matched to their relevant PDB constructs and filtered for expression system (as determined from PDB header `EXPRESSION_SYSTEM` records), discarding kinases that did not have any PDB entries with bacterial expression. As a final filtering step, the kinases were compared to three purchased kinase plasmid libraries (described in Methods), discarding kinases without a match. Construct boundaries were selected from PDB constructs and the SGC plasmid library, both of which have experimental evidence for *E. coli* expression, and subcloned from a plasmid in a purchased library (see Methods). Selecting the kinases and their constructs for this expression trial in this method rested on the basis of expected success: these specific kinase constructs were bacterially expressed and purified to a degree that a crystal structure could be solved. While expression protocols used to produce protein for crystallographic studies are often individually tailored, we considered these kinases to have a high likelihood of expressing in our semi-automated pipeline where the *same* protocol is utilized for all kinases. Statistics of the number of kinases obtained from the PDB mining procedure are shown in Figure 3A. Surprisingly, the most highly sampled family was the CAMK family, suggesting researchers may have found this family particularly amenable to bacterial expression. Based on the results of the previous experiment scanning Abl constructs for expression, we decided to use construct boundaries that were reported in the literature for each kinase. This process resulted in a set of 96 plasmid constructs distributed across kinase families (Figure 3B).

**Figure 3.**
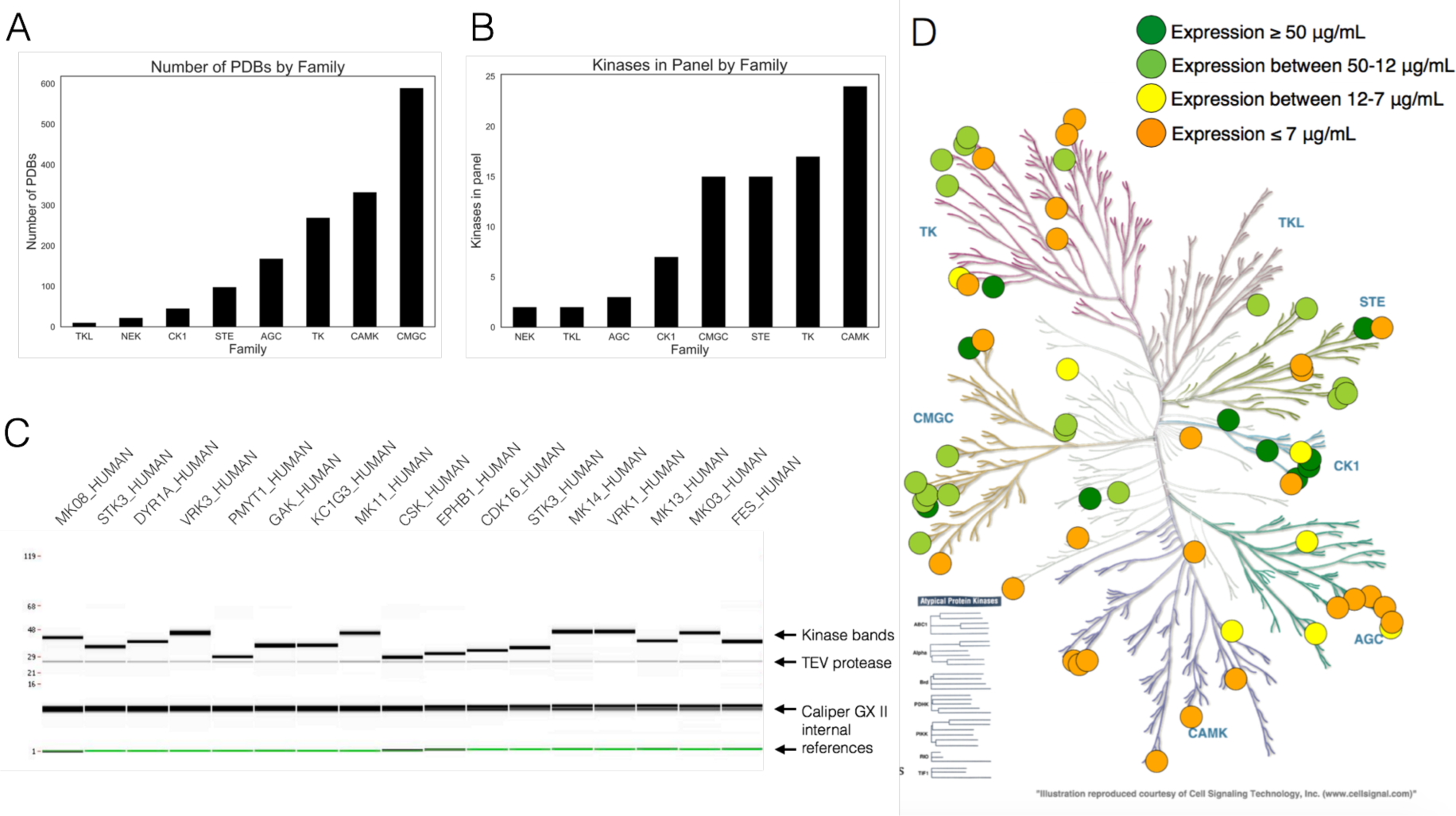
Kinome wide search for expressible kinases. (**A**) The number of PDB structures per kinase family, from the database built to select kinases for expression. (**B**) The distribution among familes of candidate kinases in our expression screen. (**C**) Caliper GX II synthetic gel image rendering of the highest expressing kinases, quantified using microfluidic capillary electrophoresis. (**D**) Kinome distribution of expression based on our 96 kinase screen. Dark green circles represent kinases with expression above 50 *µ*g/mL culture yield. Light green circles represent kinases with expression between 50 and 12 *µ*g/mL yield. Yellow circles represent kinases with expression between 12 and 7 *µ*g/mL yield. Orange circles represent kinases with any expression (even below 2 *µ*g/mL) up to 7 *µ*g/mL yield. Image made with KinMap: http://www.kinhub.org/kinmap.

From these constructs, a set of 96 His10-TEV N-terminally tagged kinase domain constructs were generated, coexpressed with a phosphatase in *E. coli*, purified via nickel bead pulldown, and quantified using microfluidic gel electrophoresis. The 96 kinases were coexpressed with either Lambda phosphatase (for Ser/Thr kinases) or a truncated form of YopH phosphatase^3^ (for Tyr kinases).

Instead of eluting with imidazole, purified kinase was cleaved off nickel beads by the addition of 10% TEV protease to minimize phosphatase contamination in the resulting eluate, allowing us to assess whether resulting yields would be sufficient (and sufficiently free of phosphatase) to permit activity assays. While the initial panel of 96 kinases was well-distributed among kinase families (Figure 3B), the most highly expressing kinases (yield of more than 12 *µ*g kinase/mL culture) were not evenly distributed (Figure 3D). While many of the kinases chosen from the CMGC and CK1 families expressed well in our panel, nearly all of the kinases from the CAMK and AGC family express below 12 *µ*g kinase/mL (Figure 3D). 52 kinases demonstrated a useful level of soluble protein expression, here defined as greater than 2 *µ*g/mL, naïvely expected to scale up to better than 2 mg/L culture (Table 1). Some kinases (shaded green in Table 1) demonstrated very high levels of expression, while others (shaded orange in Table 1) would likely benefit from further rounds of construct boundary optimization or solubility tags to boost soluble expression. The 17 most highly expressing kinases showed relatively high purity after elution, though we note that eluting via TEV site cleavage results in a quantity of TEV protease in the eluate (Figure 3C), but does not cause the elution of the His-tagged phosphatases which would hinder the ability to perform kinase activity assays. Further optimization of elution conditions may be required for optimizing kinase recovery via TEV cleavage^28-30^.

**Table 1.** Kinase domain constructs with yields >2 *µ* g/mL culture for 96-kinase expression screen. Kinases are listed by Uniprot designation and whether they were co-expressed with Lambda or truncated YopH164 phosphatase. Yield (determined by Caliper GX II quantitation of the expected size band) reported in *µ*g/mL culture, where total eluate volume was 120 *µ*L from 900 *µ*L bacterial culture. Yields are shaded green (yield > 12 *µ*g/mL), yellow (12 > yield > 7 *µ*g/mL) and orange (yield <7 *µ*g/mL); kinase domain constructs with yields that were undetectable or < 2 *µ*g/mL are not listed. ‡ denotes that the second kinase domain of KS6A1_HUMAN was expressed; all other kinases were the first or only kinase domain occurring in the ORF. Construct boundaries are listed in UniProt residue numbering for the UniProt canonical isoform. An interactive table of expression yields and corresponding constructs is available at http://choderalab.org/kinome-expression

| Kinase | Construct Boundary | Plasmid Source and ID | Phosphatase | Yield ($\mu\text{g/mL}$) |
| --- | --- | --- | --- | --- |
| MK14_HUMAN | 1-360 | Addgene 23865 | Lambda | 70.7 |
| VRK3_HUMAN | 24-352 | SGC Oxford VRK3A-c016 | Lambda | 67.5 |
| GAK_HUMAN | 24-359 | SGC Oxford GAKA-c006 | Lambda | 64.7 |
| CSK_HUMAN | 186-450 | Addgene 23941 | YopH | 62.5 |
| VRK1_HUMAN | 3-364 | Addgene 23496 | Lambda | 62.3 |
| KC1G3_HUMAN | 24-351 | SGC Oxford CSNK1G3A-c002 | Lambda | 56.3 |
| FES_HUMAN | 448-822 | Addgene 23876 | YopH | 44.0 |
| PMYT1_HUMAN | 24-311 | SGC Oxford PKMYT1A-c004 | Lambda | 38.0 |
| MK03_HUMAN | 1-379 | Addgene 23509 | Lambda | 36.4 |
| STK3_HUMAN | 16-313 | Addgene 23818 | Lambda | 34.3 |
| DYR1A_HUMAN | 24-382 | SGC Oxford DYRK1AA-c004 | Lambda | 34.1 |
| KC1G1_HUMAN | 24-331 | SGC Oxford CSNK1G1A-c013 | Lambda | 34.1 |
| MK11_HUMAN | 24-369 | SGC Oxford MAPK11A-c007 | Lambda | 31.7 |
| MK13_HUMAN | 1-352 | Addgene 23739 | Lambda | 31.7 |
| EPHB1_HUMAN | 602-896 | Addgene 23930 | YopH | 28.9 |
| MK08_HUMAN | 1-363 | HIP pJP1520 HsCD00038084 | Lambda | 28.5 |
| CDK16_HUMAN | 163-478 | Addgene 23754 | Lambda | 26.9 |
| EPHB2_HUMAN | 604-898 | HIP pJP1520 HsCD00038588 | YopH | 25.1 |
| PAK4_HUMAN | 291-591 | Addgene 23713 | Lambda | 23.9 |
| CDKL1_HUMAN | 2-304 | SGC Oxford CDKL1A-c024 | Lambda | 23.2 |
| SRC_HUMAN | 254-536 | Addgene 23934 | YopH | 22.0 |
| STK16_HUMAN | 24-316 | SGC Oxford STK16A-c002 | Lambda | 20.7 |
| MAPK3_HUMAN | 33-349 | Addgene 23790 | Lambda | 18.8 |
| PAK6_HUMAN | 383-681 | Addgene 23833 | Lambda | 18.0 |
| CSK22_HUMAN | 1-334 | HIP pJP1520 HsCD00037966 | Lambda | 17.9 |
| MERTK_HUMAN | 570-864 | Addgene 23900 | YopH | 16.8 |
| PAK7_HUMAN | 24-318 | SGC Oxford PAK5A-c011 | Lambda | 14.7 |
| CSK21_HUMAN | 1-335 | Addgene 23678 | Lambda | 14.5 |
| EPHA3_HUMAN | 606-947 | Addgene 23911 | YopH | 14.1 |
| BMPR2_HUMAN | 1-329 | SGC Oxford BMPR2A-c019 | Lambda | 14.1 |
| M3K5_HUMAN | 659-951 | HIP pJP1520 HsCD00038752 | Lambda | 14.0 |
| KCC2G_HUMAN | 24-334 | SGC Oxford CAMK2GA-c006 | Lambda | 13.3 |
| E2AK2_HUMAN | 254-551 | HIP pJP1520 HsCD00038350 | Lambda | 11.6 |
| MK01_HUMAN | 1-360 | HIP pJP1520 HsCD00038281 | Lambda | 11.2 |
| CSKP_HUMAN | 1-340 | HIP pJP1520 HsCD00038384 | Lambda | 10.1 |
| CHK2_HUMAN | 210-531 | Addgene 23843 | Lambda | 8.1 |
| KC1G2_HUMAN | 4-312 | SGC Oxford CSNK1G2A-c002 | Lambda | 7.6 |
| DMPK_HUMAN | 24-433 | SGC Oxford DMPK1A-c026 | Lambda | 7.6 |
| KCC2B_HUMAN | 11-303 | Addgene 23820 | Lambda | 7.1 |
| FGFR1_HUMAN | 456-763 | Addgene 23922 | YopH | 6.1 |
| KS6A1_HUMAN‡ | 413-735 | SGC Oxford RPS6KA1A-c036 | Lambda | 5.7 |
| DAPK3_HUMAN | 9-289 | Addgene 23436 | Lambda | 4.0 |
| STK10_HUMAN | 18-317 | HIP pJP1520 HsCD00038077 | Lambda | 3.7 |
| KC1D_HUMAN | 1-294 | Addgene 23796 | Lambda | 3.7 |
| KC1E_HUMAN | 1-294 | Addgene 23797 | Lambda | 3.5 |
| NEK1_HUMAN | 23-350 | SGC Oxford NEK1A-c011 | Lambda | 3.3 |
| CDK2_HUMAN | 1-297 | Addgene 23777 | Lambda | 3.1 |
| ABL1_HUMAN | 229-512 | HIP pJP1520 HsCD00038619 | YopH | 2.5 |
| DAPK1_HUMAN | 2-285 | HIP pJP1520 HsCD00038376 | Lambda | 2.4 |
| DYRK2_HUMAN | 23-417 | SGC Oxford DYRK2A-c023 | Lambda | 2.4 |
| HASP_HUMAN | 24-357 | SGC Oxford GSG2A-c009 | Lambda | 2.3 |
| FGFR3_HUMAN | 449-759 | Addgene 23933 | YopH | 2.3 |

Constructs with expression yields above 2 *µ*g/mL have been made available via **Addgene**: https://www.addgene.org/kits/chodera-kinase-domains

### High-expressing kinases are folded with a well-formed ATP binding site

To determine whether the expressed kinases were properly folded, we performed both a fluorescence-based thermostability assay (Figure 4) as well as a fluorescent ATP-competitive ligand binding measurement to quantify whether the ATP binding site was well-formed (Figure 5).

**Figure 4.**
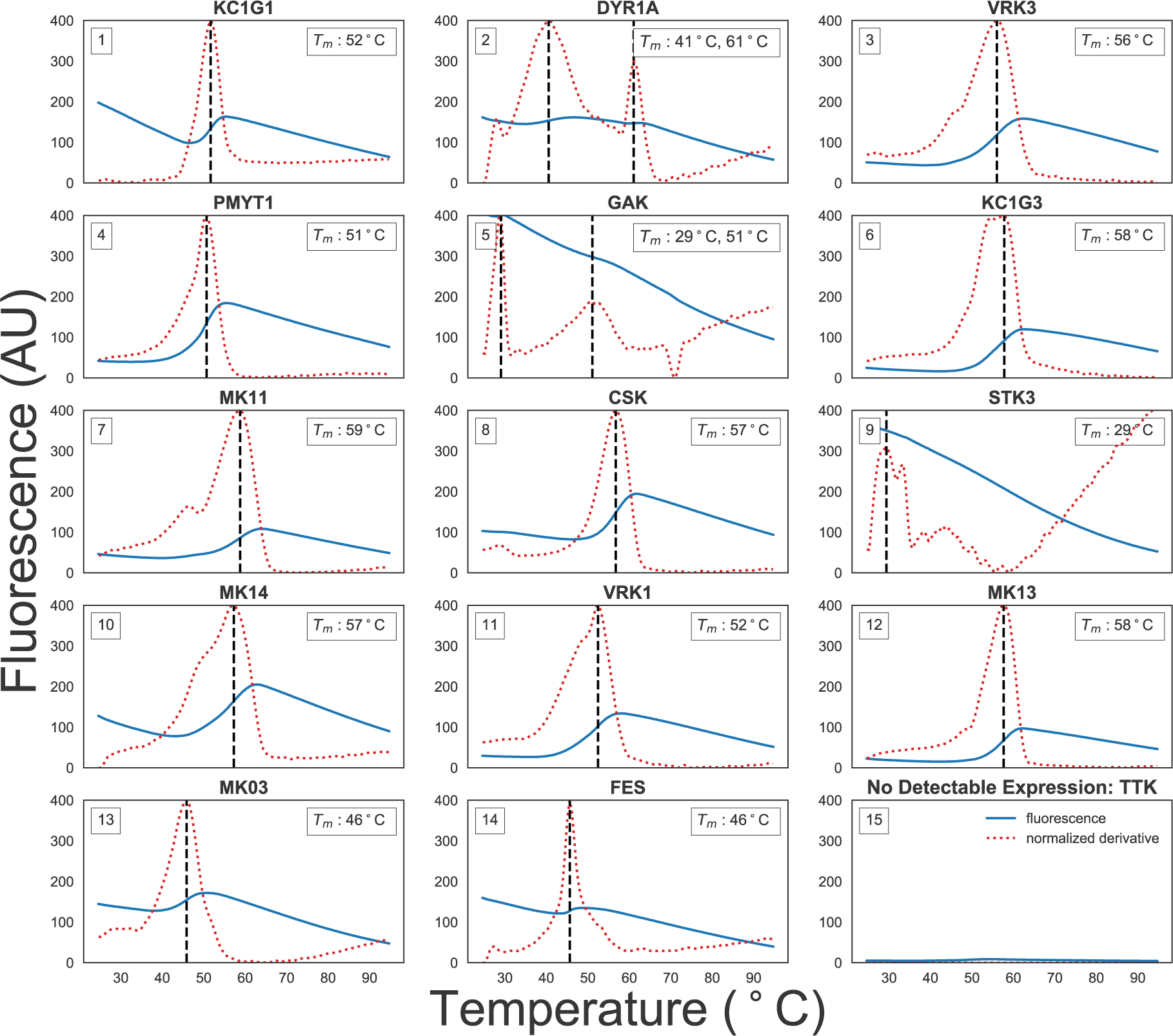
Fluorescence-based thermostability assay demonstrates many high-expressing kinases are well-folded. A fluorescence-based thermostability assay was performed on the 14 kinases shown to express above a minimum 0.24 mg/mL concentration after elution. SYPRO Orange fluorescence (solid blue line) was measured at 580 nm (half bandwidth 20 nm) after excitation at 465 nm (half bandwith 25 nm) as as the temperature was ramped from (x-axis) in Nickel Buffer A (25 mM HEPES pH 7.5, 5% glycerol, 400 mM NaCl, 20 mM imidazole, 1 mM BME). The temperature was held at 25°C for 15 sec before ramping up to 95°C with a ramp rate of 0.06°C/s. The unfolding temperature *T*_*m*_ (black dashed line and insert) was determined from the maxima of the normalized first derivative of fluorescence (red dashed line). Fluorescence emission at 580 nm is shown on the left y-axis. To control for signals resulting from TEV protease contamination present at 0.01-0.03 mg/mL, TTK, a kinase with no detectable expression in our panel as determined via Caliper GX II quantitation was in included (panel 15).

**Figure 5.**
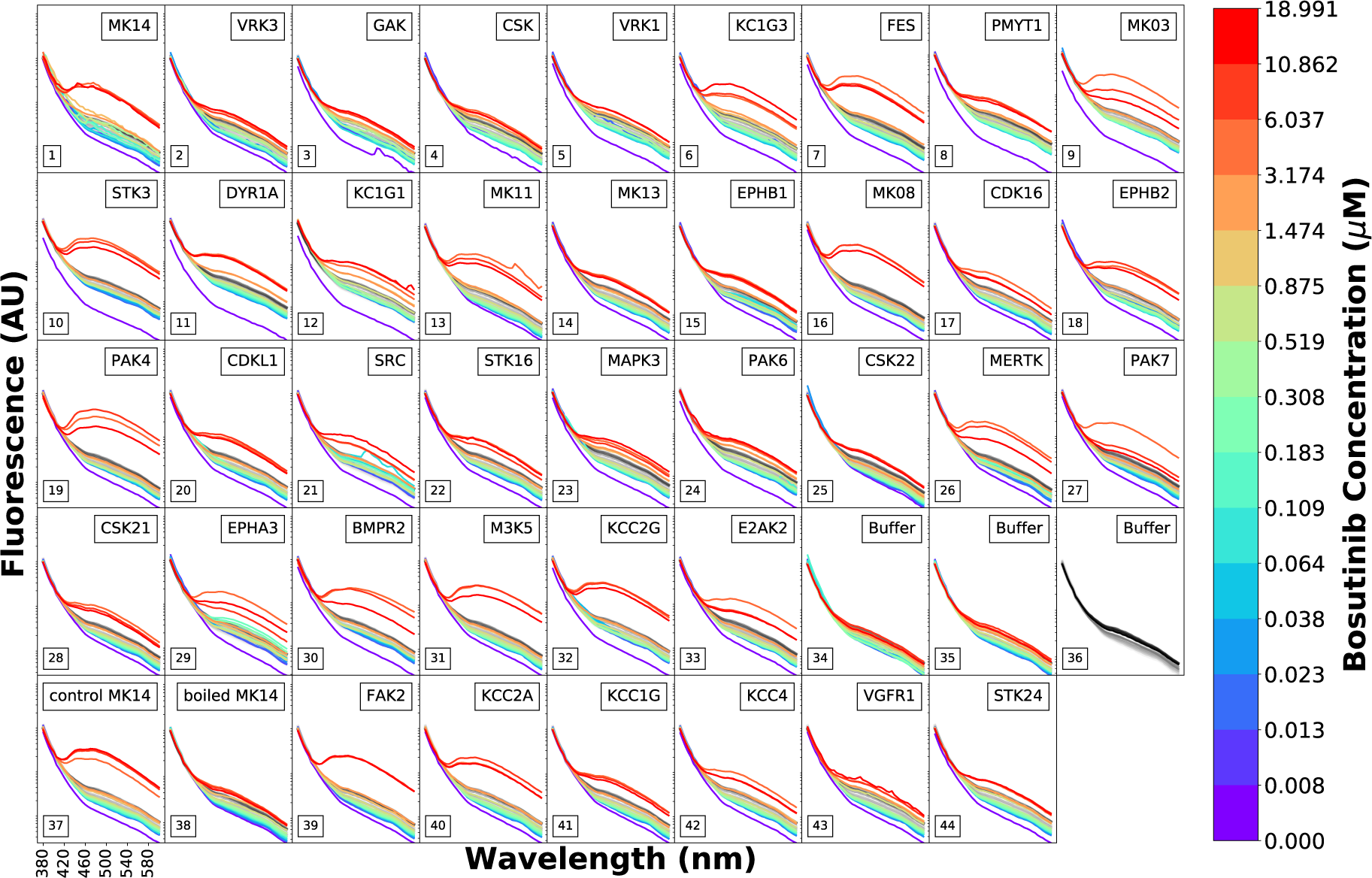
Fluorescence emission spectra as a function of the fluorescent ATP-competitive kinase inhibitor bosutinib demonstrates the presence of a well-formed ATP binding pocket. The ATP-competitive inhibitor bosutinib shows a strong increase in fluorescence centered around 450 nm when bound to kinases with well-folded ATP binding sites upon excitation at 280 nm^34^. To assess whether the kinases from the high-throughput expression screen were well-folded, bosutinib was titrated in a 15-concentration series geometrically spanning 0.008 *µ*M to 18.99 *µ*M (colored lines, higher concentrations are shown in warmer colors) in 15 increments for 39 expressing kinases with protein concentration adjusted to ~0.5 *µ*M in 100 *µ*L assay volume. Eluted TEV protease contaminant varies from 0.01-0.03 mg/mL in the assay volumes. The control MK14 and boiled MK14 (boiled for 10 min at 95°C) were produced in a large scale expression from the same plasmid as used in the high-throughput expression protocol and they were included as positive and negative controls for bosutinib binding to ATP binding pocket. Fluorescence emission spectra (y-axis, bandwidth 20 nm) were measured from 370 nm to 600 nm (x-axis) for excitation at 280 nm (bandwidth 10 nm). For reference, the fluorescence of bosutinib titrated into buffer titration (panel 36) is shown in grayscale in each panel. Significant increases in fluorescence signal above baseline qualitatively indicate the presence of a well-formed ATP binding site.

#### Fluorescence-based thermostability assay

A fluorescence-based thermostability assay was performed with the hydrophobic dye SYPRO Orange to determine whether a strong two-state unfolding signal could be observed (see Methods). Also referred to as *thermofluor* or *differential scanning fluorimetry (DSF)*, as the temperature is slowly increased, unfolded proteins will expose hydrophobic patches that SYRPO orange will bind to, causing an increase in fluorescence^31-33^. While the fluorescence of solvated SYPRO Orange is temperature-dependent, clear unfolding temperatures (*T*_*m*_) can often be identified from peaks in the first derivative of the observed fluorescence signal. Figure 4 shows the fluorescence (blue line), the absolute value of its derivative (red dashed line), and the unfolding temperature determined from the maximum absolute derivative (*T*_*m*_) for the the 14 kinases that were eluted to concentrations above 0.24 mg/mL eluate, which was determined to be the minimum concentration required for optimal resolution of melting curves upon dilution to 10 *µ*L. Because TEV-eluted kinase was used directly in this assay, TEV protease contaminant varies from 0.01-0.03 mg/mL in the resulting assay mix. The selected minimum concentration ensured that the kinase was roughly an order of magnitude higher concentration than the contaminating TEV.

Most of the kinases assayed had strong peaks above room temperature, suggesting that they are well-folded in the elution buffer (25 mM HEPES pH 7.5, 5% glycerol, 400 mM NaCl, 20 mM imidazole, 1 mM BME) at room temperature. Some kinases, such as a DYR1A and GAK (Figure 4, panels 6 and 9), had two shallow inflection points in SYPRO fluorescence as a function of temperature. While STK3 does not have a strong peak above room temperature, titration with an ATP-competitive inhibitor suggests this kinase either has a well-formed ATP binding site or folding can be induced by ligand binding (Figure 5, panel 10). As a control, a sample with no detectable kinase expression (TTK from our expression panel) was assayed (Figure 4, panel 9), which showed nearly no fluorescence signal.

#### ATP-competitive inhibitor binding fluorescence assay

To determine whether expressed kinases had well-folded ATP binding sites, we probed their ability to bind an ATP-competitive inhibitor. While a pan-kinase inhibitor such as staurosporine could be used as a fluorescent probe^35^, the ATP-competitive inhibitor bosutinib shows a much stronger increase in fluorescence around 450-480 nm when bound to kinases with well-folded ATP binding sites^34, 36^. While excitation at 350 nm can be used, excitation at 280 nm results in lower background, potentially due to fluorescent energy transfer between kinase and ligand. Despite the weak affinity of bosutinib for many kinases, its aqueous solubility is sufficient to provide a quantitative assessment of ATP-competitive binding to many kinases at sufficiently high concentrations to function as a useful probe^34, 36^.

Here, we utilized this approach as a *qualitative* probe for ATP-competitive ligand binding, due to uncertainty in the ligand concentration caused by significant evaporation over the course of the sequential titration experiment (see Methods section for a more in depth discussion). 33 of the kinases in our expression panel had sufficient yields to prepare 100 *µ*L of 0.5 *µ*M kinase assay solutions, and were assessed for binding to bosutinib (Figure 5, panels 1-33), with a concentration-dependent increase in fluorescence signal (colored spectra) over the baseline ligand fluorescence titrated into buffer (gray spectra) providing evidence of a well-formed ATP binding site. Six of the lowest expression kinase constructs (Figure 5, panels 39-44) were prepared by diluted 20 *µ*L to a reaction volume of 100 *µ*L and assessed for bosutinib binding. Unexpectedly, these kinases also showed evidence of binding, suggesting this assay is able to detect a well-formed ATP binding site even for protein concentrations less than 0.5 *µ*M. To demonstrate that unfolded kinases do not demonstrate this increase in fluorescence over ligand-only baseline, thermally denatured MK14 was included as a control next to folded MK14 from a large-scale expression prep (Figure 5, panels 37–38), with thermally denatured MK14 exhibiting little difference from titrating ligand into buffer alone.

### Expressing clinically-derived Src and Abl mutants

Next-generation sequencing has enabled generation of massive datasets rich with missense alterations in kinases observed directly in the clinic^37-39^, and has been particularly transformative in the field of oncology. To determine how well our human kinase domain panel supports the automated expression of clinically-identified missense mutants for biophysical, biochemical, and structural characterization, we attempted to express 96 missense mutations mined from sequencing studies of cancer patients. The mutations were gathered using cBioPortal^40^ from publicly available sources and a large clinical tumor sequencing dataset from the Memorial Sloan Kettering Cancer Center^38^ sequenced in the MSK-IMPACT panel^41^.

Using our structural informatics pipeline, a database was built focusing on the kinases we found to be expressible in *E. coli*. To add the mutation data, we retrieved public datasets from cBioPortal^44, 45^ along with annotations from Oncotator^46^ through their respective web service APIs. We then added mutations and annotations from the MSKCC dataset^38^ by extracting the mutations from a local copy of the dataset and retrieving annotations from Oncotator. The annotated mutations were filtered for mutations that occurred within the construct boundaries of our kinase domains. We found 63 unique clinical mutations appearing within our kinase domain construct boundaries for Abl and 61 for Src. We subsequently selected 48 mutants for Abl and 46 for Src to express, aiming for a panel of mutants distributed throughout the kinase domain (Figure 6A), with wild-type sequences included as controls. Mutations were introduced using site-directed mutagenesis and assayed for expression yields (Figure 6B). Those with yields above 2 *µ*g kinase/mL culture are listed in Table 2.

**Figure 6.**
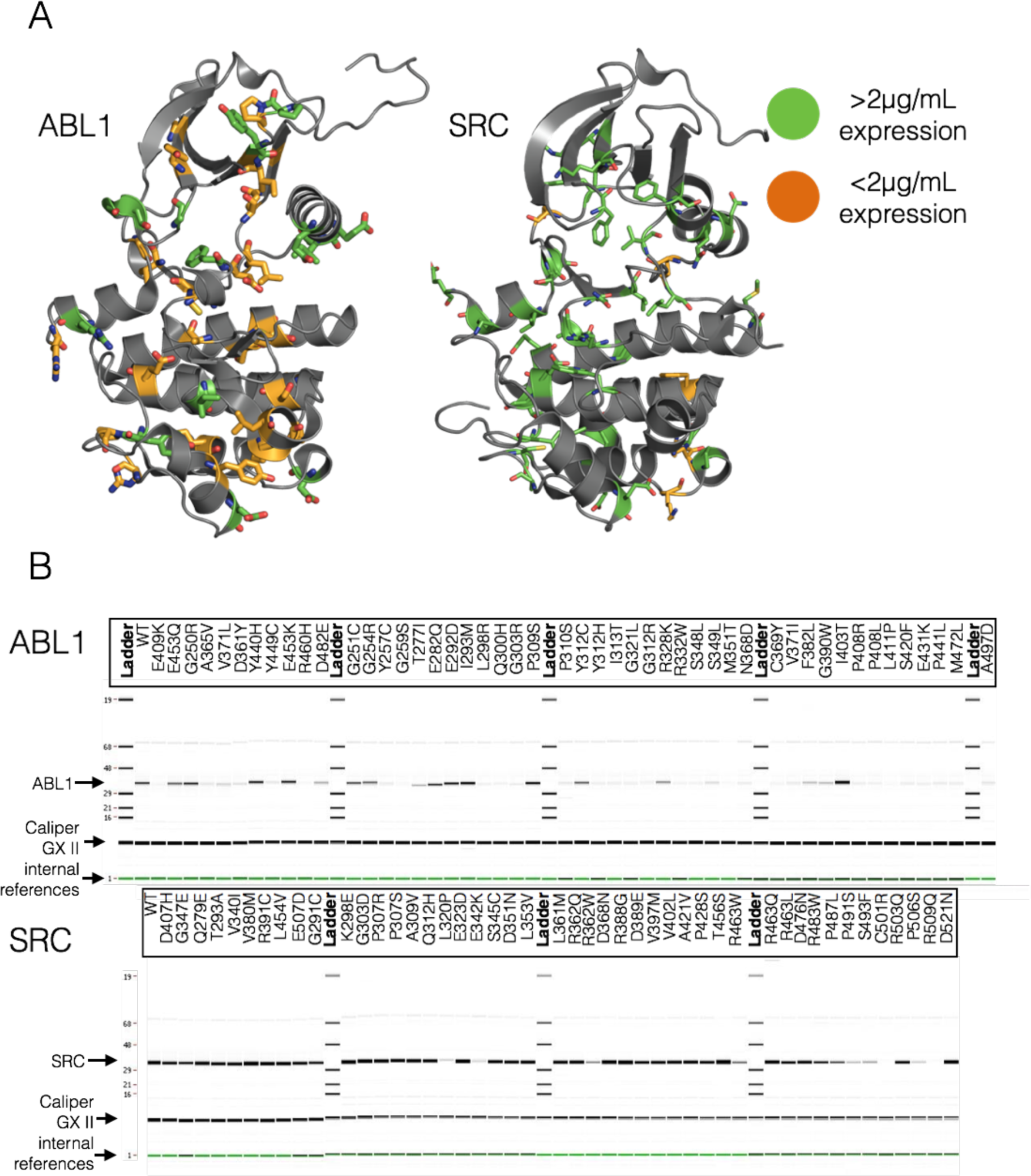
Expression yields for engineered clinically-derived Src and Abl missense mutants. (**A**) All Abl and Src clinically-identified mutants assessed in the expression screen are displayed as sticks. Mutants with expression yields >2 *µ*g/mL are colored green, while those with yields <2 *µ*g/mL are colored orange. Rendered structures are Abl (PDBID: 2E2B) and Src (PDBID: 4MXO) 36. (**B**) Synthetic gel images showing ABl (*top*) or Src (*bottom*) expression, with wells labeled by missense mutation. Yield was determined by Caliper GX II quantitation of the expected size band and reported in *µ*g/mL culture, where total eluate volume was 120 *µ*L following nickel bead pulldown purification from 900 *µ*L bacterial culture. Residue mutations use numbering for the Uniprot canonical isoform.

**Table 2.** Expression yields for engineered clinical missense mutants of Src and Abl kinase domains with yields > 2 *µ* g/mL culture. Src and Abl kinase domain constructs with engineered clinical mutations with expression yields >2 *µ*g/mL culture are listed, sorted by yield. Yield was determined by Caliper GX II quantitation of the expected size band and reported in *µ*g/mL culture, where total eluate volume was 80 *µ*L purified from 900 *µ*L bacterial culture. Wild-type (WT) controls for both Src and Abl (here, a single well for each) are shown as the first entry for each gene.

| Abl1 (229-512) | Mutation<sup>1</sup> | Functional Impact Score<sup>2</sup> | yield ($\mu\text{g/mL}$) | % of WT expression |
| --- | --- | --- | --- | --- |
|  | WT | – | 5.1 | – |
|  | I403T | Low | 17.8 | 350 |
|  | I293M | Low | 9.8 | 193 |
|  | P309S | Neutral | 7.8 | 153 |
|  | E453K | Low | 7.3 | 144 |
|  | Y440H | Medium | 7.1 | 140 |
|  | E292D | Low | 6.9 | 135 |
|  | G251C | High | 5.2 | 102 |
|  | E282Q | Neutral | 5.1 | 102 |
|  | G250R | Neutral | 5.1 | 100 |
|  | G254R | High | 5.0 | 98 |
|  | Y312C | Neutral | 4.7 | 93 |
|  | E453Q | Low | 3.7 | 73 |
|  | R328K | Low | 3.5 | 69 |
|  | D482E | Neutral | 2.5 | 49 |
|  | F382L | Medium | 2.1 | 41 |
|  | G390W | Medium | 2.1 | 41 |
| Src (254-536) | Mutation<sup>1</sup> | Functional Impact Score<sup>2</sup> | yield ($\mu\text{g/mL}$) | % of WT expression |
|  | WT | – | 35.7 | – |
|  | T456S | Neutral | 80.9 | 227 |
|  | R388G | Medium | 61.5 | 172 |
|  | K298E | High | 54.5 | 153 |
|  | V380M | Neutral | 51.7 | 145 |
|  | D368N | Neutral | 49.9 | 140 |
|  | D521N | Low | 42.8 | 120 |
|  | R463Q | Neutral | 38.4 | 108 |
|  | R391C | Neutral | 37.5 | 105 |
|  | E323D | Low | 37.2 | 104 |
|  | A309V | Low | 35.9 | 98 |
|  | G303D | Neutral | 34.1 | 96 |
|  | R362Q | Neutral | 33.6 | 94 |
|  | L361M | Medium | 31.7 | 89 |
|  | A421V | Neutral | 30.7 | 86 |
|  | V402L | Neutral | 30.6 | 86 |
|  | V397M | Medium | 29.8 | 84 |
|  | Q278E | Neutral | 29.6 | 83 |
|  | Q312H | Low | 29.5 | 83 |
|  | L353V | Medium | 29.0 | 81 |
|  | L454V | Neutral | 29.0 | 81 |
|  | P307R | Neutral | 28.6 | 80 |
|  | V340I | Low | 28.0 | 78 |
|  | P307S | Neutral | 24.2 | 68 |
|  | D476N | Neutral | 23.3 | 65 |
|  | D351N | Neutral | 22.9 | 64 |
|  | T293A | Neutral | 22.2 | 62 |
|  | S345C | Low | 22.2 | 62 |
|  | P428S | Medium | 22.2 | 62 |
|  | E507D | Neutral | 20.7 | 58 |
|  | D389E | High | 20.0 | 56 |
|  | R503Q | Neutral | 17.3 | 49 |
|  | D407H | High | 15.9 | 45 |
|  | R463L | Neutral | 14.9 | 42 |
|  | G291C | Medium | 11.9 | 33 |
|  | G347E | Medium | 10.2 | 29 |
|  | R483W | High | 9.8 | 27 |
|  | P487L | Medium | 6.0 | 17 |
|  | R463W | Medium | 5.2 | 15 |
|  | R362W | Low | 3.9 | 11 |
|  | S493F | Low | 3.0 | 8 |
|  | P491S | Low | 2.2 | 6 |

<sup>1</sup> Uniprot amino acid sequence numbering of primary isoform
<sup>2</sup> MutationAssessor Score<sup>42,43</sup>, which predicts functional impact via conservation

High-expressing mutants appear to be distributed relatively uniformly throughout the kinase domain (Figure 6A). While the vast majority of the Src mutants expressed at a usable level, many of the Abl mutants expressed below the 2 *µ*g/mL threshold. This can primarily be attributed to the low level of expression for wild-type Abl construct (Table 1). In instances where kinase activity is not required, yield could be increased via the introduction of inactivating mutations 21 or further tailoring of expression and purification protocols.

## Methods

### Semi-automated selection of kinase construct sequences for *E. coli* expression

#### Selection of human protein kinase domain targets

Human protein kinases were selected by querying the UniProt API (query date 30 May 2014) for any human protein with a domain containing the string “protein kinase”, and which was manually annotated and reviewed (i.e. a Swiss-Prot entry). The query string used was:

`taxonomy:"Homo sapiens (Human) [9606]" AND domain:"protein kinase" AND reviewed:yes`

Data was returned by the UniProt API in XML format and contained protein sequences and relevant PDB structures, along with many other types of genomic and functional information. To select active protein kinase domains, the UniProt domain annotations were searched using the regular expression `ˆProtein kinase(?!; truncated)(?!; inactive)`, which excludes certain domains annotated “Protein kinase; truncated” and “Protein kinase; inactive”. Sequences for the selected domains, derived from the canonical isoform as determined by UniProt, were then stored.

#### Matching target sequences with relevant PDB constructs

Each target kinase gene was matched with the homologous in any other species, if present, and all UniProt data was downloaded. This data included a list of PDB structures which contain the protein, and their sequence spans in the coordinates of the UniProt canonical isoform. PDB structures which did not include the protein kinase domain or truncated more than 30 residues at each end were filtered out. PDB coordinate files were then downloaded for each remaining PDB entry. The coordinate files contain various metadata, including the `EXPRESSION_SYSTEM` annotation, which was used to filter PDB entries for those which include the phrase “ESCHERICHIA COLI”. The majority of PDB entries returned had an `EXPRESSION_SYSTEM` tag of “ESCHERICHIA COLI”, while a small number had “ESCHERICHIA COLI BL21” or “ESCHERICHIA COLI BL21(DE3)”.

The PDB coordinate files also contain SEQRES records, which should contain the protein sequence used in the crystallography or NMR experiment. According to the PDB File Format FAQ (http://deposit.rcsb.org/format-faq-v1.html), "All residues in the crystal or in solution, including residues not present in the model (i.e., disordered, lacking electron density, cloning artifacts, HIS tags) are included in the SEQRES records." However, we found that these records are very often misannotated, instead representing only the crystallographically resolved residues. Since expression levels can be greatly affected by insertions or deletions of only one or a few residues at either terminus^47^, it is important to know the full experimental sequence. To measure the authenticity of a given SEQRES record, we developed a simple metric by hypothesizing that most crystal structures would likely have at least one or more unresolved residues at one or both termini and that the presence of an expression tag, which is typically not crystallographically resolved, would indicate an authentic SEQRES record. To achieve this, unresolved residues were first defined by comparing the SEQRES sequence to the resolved sequence, using the SIFTS service to determine which residues were not present in the canonical isoform sequence^48^. Regular expression pattern matching was used to detect common expression tags at the N- or C-termini. Sequences with a detected expression tag were given a score of 2, while those with any unresolved sequence at the termini were given a score of 1, and the remainder were given a score of 0. This data was stored to allow for subsequent selection of PDB constructs based on likely authenticity in later steps. The number of residues extraneous to the target kinase domain, and the number of residue conflicts with the UniProt canonical isoform within that domain span were also stored for each PDB sequence.

#### Plasmid libraries

As a source of kinase DNA sequences for subcloning, we purchased three kinase plasmid libraries: the Addgene Human Kinase ORF kit, a kinase library from the Structural Genomics Consortium (SGC), Oxford (http://www.thesgc.org), and a kinase library from the PlasmID Repository maintained by the Dana-Farber/Harvard Cancer Center. Annotated data for the kinases in each library was used to match them to the human protein kinases selected for this project. The plasmid open reading frames (ORFs) were translated into protein sequences and aligned against the target kinase domain sequences from UniProt. Also calculated were the number of extraneous protein residues in the ORF, relative to the target kinase domain sequence, and the number of residue conflicts with the UniProt sequence. Our aim was to subclone the chosen sequence constructs from these library plasmids into our expression plasmids.

#### Selection of sequence constructs for expression

Of the kinase domain targets selected from UniProt, we filtered out those with no matching plasmids in our available plasmid libraries or no suitable PDB construct sequences. For this purpose, a suitable PDB construct sequence was defined as any with an authenticity score greater than zero (see above). Library plasmid sequences and PDB constructs were aligned against each Uniprot target domain sequence, and various approaches were considered for selecting the construct boundaries to use for each target, and the library plasmid to subclone it from. Candidate construct boundaries were drawn from two sources: PDB constructs and the SGC plasmid library, has been successfully tested for *E. coli* expression.

For most of the kinase domain targets, multiple candidate construct boundaries were available. To select the most appropriate construct boundaries, we sorted them first by authenticity score, then by the number of conflicts relative to the UniProt domain sequence, then by the number of residues extraneous to the UniProt domain sequence span. The top-ranked construct was then chosen. In cases where multiple library plasmids were available, these were sorted first by the number of conflicts relative to the UniProt domain sequence, then by the number of residues extraneous to the UniProt domain sequence span, and the top-ranked plasmid was chosen. This process resulted in a set of 96 kinase domain constructs, which (by serendipity) matched the 96-well plate format we planned to use for parallel expression testing. We selected these constructs for expression testing.

An interactive table of the selected plasmids, constructs, and aligned PDB files can be viewed at http://choderalab.org/kinome-expression.

#### Automation of the construct selection process

While much of this process was performed programmatically, many steps required manual supervision and intervention to correct for exceptional cases. While these exceptions were encoded programmatically as overrides to ensure the scheme could be reproduced from existing data, we hope to eventually develop a fully automated software package for the selection of expression construct sequences for a given protein family, but this was not possible within the scope of this work.

#### Mutagenesis protocol

Point mutations were introduced with a single-primer QuikChange reaction. Primers were designed to anneal at 55°C both upstream and downstream of the point mutation, and with a total length of approximately 40 bases. At the codon to be modified, the fewest possible number of bases was changed. Plasmid template (160 ng) was mixed with 1 *µ*M primer in 1x PfuUltra reaction buffer, with 0.8 mM dNTPs (0.2 mM each) and 1 U PfuUltra High-Fidelity DNA polymerase (Agilent), in a total volume of 20 *µ*L. Thermocycler settings were 2 min at 95°C, followed by 18 cycles of 20s at 95°C, 1 min at 53°C, 12 min at 68°C (2min/kb), then 1 minute at 68°C. After cooling to room temperature, 4 *µ*L of the PCR reaction was added to 16 *µ*L CutSmart Buffer (NEB) containing 10 U DpnI (NEB). After incubation for 2.5 hours at 37°C, 6 *µ*L of this mixture was used to directly transform XL1-Blue chemically competent cells (Agilent) according to the manufacturer’s protocol. Transformants were picked for plasmid mini-preps and the presence of the point mutations was confirmed by sequencing.

#### Expression testing

For each target, the selected construct sequence was subcloned from the selected DNA plasmid. Expression testing was performed at the QB3 MacroLab (QB3 MacroLab, University of California, Berkeley, CA 94720) [http://qb3.berkeley.edu/macrolab/], a core facility offering automated gene cloning and recombinant protein expression and purification services.

Each kinase domain was tagged with a N-terminal His10-TEV and coexpressed with either the truncated YopH164 for Tyr kinases or lambda phosphatase for Ser/Thr kinases. All construct sequences were cloned into the 2BT10 plasmid, an AMP resistant ColE1 plasmid with a T7 promoter, using ligation-independent cloning (LIC). The inserts were generated by PCR using the LICv1 forward (TACTTCCAATCCAATGCA) and reverse (TTATCCACTTCCAATGTTATTA) tags on the primers. Gel purified PCR products were LIC treated with dCTP. Plasmid was linearized, gel purified, and LIC-treated with dGTP. LIC-treated plasmid and insert were mixed together and transformed into XL1-Blues for plasmid preps.

Expression was performed in Rosetta2 cells (Novagen) grown with Magic Media (Invitrogen autoinducing medium), 100 *µ*g/mL of carbenicillin and 100 *µ*g/mL of spectinomycin. Single colonies of transformants were cultivated with 900 *µ*L of MagicMedia into a gas permeable sealed 96-well block. The cultures were incubated at 37°C for 4 hours and then at 16°C for 40 hours while shaking. Next, cells were centrifuged and the pellets were frozen at −80°C overnight. Cells were lysed on a rotating platform at room temperature for an hour using 700 *µ*L of SoluLyse (Genlantis) supplemented with 400 mM NaCl, 20 mM imidazole, 1 *µ*g/mL pepstatin, 1 *µ*g/mL leupeptin and 0.5 mM PMSF.

For protein purification, 500 *µ*L of the soluble lysate was added to a 25 *µ*L Protino Ni-NTA (Machery-Nagel) agarose resin in a 96-well filter plate. Nickel Buffer A (25 mM HEPES pH 7.5, 5% glycerol, 400 mM NaCl, 20 mM imidazole, 1 mM BME) was added and the plate was shaken for 30 min at room temperature. The resin was washed with 2 mL of Nickel Buffer A. For the 96-kinase expression experiment, target proteins were eluted by a 2 hour incubation at room temperature with 10 *µ*g of TEV protease in 80 *µ*L of Nickel Buffer A per well and a subsequent wash with 40 *µ*L of Nickel Buffer A to maximize protein release. Nickel Buffer B (25 mM HEPES pH 7.5, 5% glycerol, 400 mM NaCl, 400 mM imidazole, 1 mM BME) was used to elute TEV resistant material remaining on the resin. Untagged protein eluted with TEV protease was run on a LabChip GX II Microfluidic system to analyze the major protein species present.

For the clinical mutant and Abl1 construct boundaries expression experiments, target proteins were washed three times with Nickel Buffer A prior to elution in 80 *µ*L Nickel Buffer B. The eluted protein was run on a LabChip GX II Microfluidic system to analyze with major protein species were present.

#### Fluorescence-based thermostability assay

To assess whether the highly-expressed wild-type kinase constructs are folded, a thermofluor thermostability assay 31-33 was performed for kinase constructs that have a minimum of 0.24 mg/mL protein concentration in the eluate. After diluting 9 *µ*L of eluate by 1 *µ*L dye, the effective assay concentration is 0.216 mg/mL minimum in 10 *u*L assay volume. Previous optimization efforts in the lab determined that 0.20 mg/mL was the lower limit of well-defined T_*m*_ detection. This minimum concentration also ensured that the kinase was present at roughly an order of magnitude concentration higher than contaminating TEV protease.

Kinase expression panel eluates, which were kept in 96-well deep well plate frozen at −80°C for 2 years prior to the thermal stability assay, were thawed in an ice-water bath for 30 min. 9 *µ*L of each kinase eluate was added to a 384 well PCR plate (4titude-0381). 100X SYPRO Orange dye solution was prepared from a 5000X DMSO solution of SYPRO Orange Protein Gel Stain (Life Technologies, Ref S6650, LOT 1790705) by dilution in distilled water. In initial experiments, SYPRO Orange dye solution was diluted in kinase binding assay buffer (20 mM Tris 0.5 mM TCEP pH 8), which caused the dye to precipitate out of solution. Particulates in the dye solution were pelleted by tabletop centrifugation (2 min, 5000 RCF) and the solution was kept covered with aluminum foil in the dark to prevent photodamage. 1 *µ*L of 100X dye solution was added to each kinase eluate sample in 384-well PCR plate. The plate was sealed with Axygen UC-500 Ultra Clear Pressure Sensitive sealing film. To remove any air bubbles, the sample plate was centrifuged for 30 sec with 250 g using Bionex HiG4 centrifuge. Sample mixing was performed by orbital shaking with Inheco shakers for 2 min at 1000 RPM.

A thermofluor melt was performed using a LightCycler 480 (Roche) qPCR instrument using an excitation filter of 465 nm (half bandwidth 25 nm) and emission filter at 580 nm (half bandwidth 20 nm). LightCycler 480 Software Version 1.5.1 was used to operate the instrument and analyze the results. The temperature was held at 25°C for 15 s before ramping up to 95°C with a ramp rate of 0.06°C/s. During temperature ramp 10 fluorescence acquisitions/°C were recorded with dynamic integration time mode, melt factor of 1, quant factor of 10, and maximum integration time of 2 sec. Thermal protein denaturation causes hydrophobic patches of protein to be exposed, which SYPRO Orange dye can bind. Binding of SYPRO Orange dye is detected as an increase in fluorescence at 580 nm. Presence of a clear thermal denaturation peak in the absolute value of the derivative of the fluorescence as a function of temperature serves as an indication that the proteins were well-folded. Observed fluorescence was plotted as a function of temperature, and a melting temperature *T*_*m*_ was determined as the maximum of the absolute value of its first derivative.

#### ATP-competitive inhibitor binding fluorescence assay

To determine whether the expressed kinases had a well-folded ATP-binding site, we assessed whether the eluted kinase was capable of binding the ATP-competitive small molecule kinase inhibitor bosutinib. We designed fluorescence-based binding assays following earlier work reporting that this quinoline-scaffold inhibitor undergoes a strong increase in fluorescence upon binding (even weakly) to kinase ATP-binding sites^34^. By titrating in the ligand to close to the solubility limit, even weak binding to the ATP-binding site can be detected by observing emission increases around 450 nm during excitation at 280 nm.

For 33 of the kinases in our expression panel, 0.5 *µ*M kinase solutions from kinase expression panel eluates were prepared in kinase binding assay buffer (20 mM Tris 0.5 mM TCEP pH 8) for a final volume of 100 *µ*L in a black 96-well vision plate (4titude-0223). Six low-expressing kinases (Figure 5, panels 39-44) were prepared by diluting 20 *µ*L of eluate in kinase binding assay buffer (20 mM Tris 0.5 mM TCEP pH 8) to a final volume of 100 *µ*L, for final concentrations below 0.5 *µ*M. The plate was shaken for 2 min clockwise and 2 min counter-clockwise by orbital shaking with Inheco shakers at 2000 RPM and centrifuged for 30 sec with 1000 g using Bionex HiG4 centrifuge. Fluorescence emission spectra were measured from 370 nm to 600 nm (20 nm bandwidth) in 5 nm steps using 280 nm excitation (10 nm bandwidth) from both the top and bottom of the well using a Tecan Infinite M1000 PRO.

Bosutinib free base (LC Labs, cat no. B-1788, lot no. BSB-103, M.W. 530.45 Da) was dispensed directly from a roughly 10 mM DMSO stock solution to the assay solution using a Tecan HP D300 Digital Dispenser. The 10 mM DMSO stock solution was prepared gravimetrically using an automated balance (Mettler Toledo Balance XPE205 with LabX Laboratory Software) by dispensing 39.02 mg solid Bosutinib powder stored under nitrogen gas at 25°C into 8.0499 g DMSO (Alfa Aesar, cat no. 42780, log no. Y25B604, density 1.1004 g/mL at ambient temperature) which is kept dry under argon gas at 25°C. To minimize atmospheric water absorption due to the hygroscopic nature of DMSO, the 10 mM stock solution was pipetted into wells of a 96-well stock plate by an automated liquid handling device (Tecan EVO 200 with air LiHa) and sealed with foil seal (PlateLoc). Ligand was dispensed to the assay plate with HP D300 (using aliquots of stock solution pipetted from a freshly pierced stock plate well) targeting a roughly geometrically-increasing series of ligand concentrations in each well to achieve the following total ligand concentrations after each dispense: 0.008 *µ*M, 0.013 *µ*M, 0.023 *µ*M, 0.038 *µ*M, 0.064 *µ*M, 0.109 *µ*M, 0.183 *µ*M, 0.308 *µ*M, 0.519 *µ*M, 0.875 *µ*M, 1.474 *µ*M, 3.174 *µ*M, 6.037 *µ*M, 10.862 *µ*M, 18.991 *µ*M. The plate was shaken by HP D300 for 10 sec after usage of each dispensehead. After each titration, the plate was shaken with Inheco shakers (2 min clockwise and counter-clockwise, 2000 RPM, orbital shaking) and centrifuged (30 sec, 1000 g) using a Bionex HiG4 centrifuge. Fluorescence spectra from 370 nm to 600 nm (bandwith 20 nm) in 5 nm steps using 280 nm excitation (bandwidth 10 nm) were read from both the top and bottom of the well using a Tecan Infinite M1000 PRO. In total, the experiment took 17.5 hours to complete due to the time-consuming spectral read after each dispense, likely resulting in significant evaporation from some wells during the experiment.

ATP-competitive binding was analyzed qualitatively for each kinase by plotting the fluorescence spectra as a function of concentration to detect concentration-dependent increases in fluorescence. As a control for background ligand fluorescence independent of protein binding, fluorescence spectra of three replicates of ligand into buffer titrations were plotted. As a positive control, MK14 produced by a validated large scale expression protocol (see Supplementary Methods) from the same plasmid used in the high-throughput protocol was included. To control for non-specific binding to unfolded protein, we included boiled MK14 (prepared from the large scale expression of MK14 by boiling at 95°C for 10 min). A concentration-dependent increase in fluorescence was interpreted as evidence that the ATP-binding site of the kinase was well folded and allowed for bosutinib binding. Due to the length of the experiment, it is possible that evaporation reduced the well volume below 100 *µ*L and potentially caused bosutinib to reach higher concentration levels than expected. This creates uncertainty for data points, as bosutinib may either be a higher concentration (due to evaporation) or a lower concentration (due to potential precipitation caused by lower well volumes) than expected. For this reason, we have interpreted the experiment as qualitative evidence of binding, instead of quantitatively. Bosutinib binding is an indication of proper folding of the ATP binding pocket of these recombinantly expressed kinase constructs.

## Discussion

We have demonstrated that a simple, uniform, automatable protocol is able to achieve useful bacterial expression yields for a variety of kinase domain constructs. While yields could likely be further improved by a variety of methods-such as the addition of solubility—promoting tags, construct domain boundary and codon optimization, or mutations to improve the solubility or ablate catalytic activity—the simplicity of this approach suggests widespread utility of automated bacterial expression for biophysical, biochemical, and structural biology work for the further study of human kinase domains.

Our expression test of 81 different construct boundaries of the Abl kinase domain demonstrated a surprising sensitivity of expression yields to the precise choice of boundary. This sensitivity may be related to where the construct is truncated with respect to the secondary structure of the protein, as disrupting secondary structure could cause the protein to improperly fold, leading to low soluble protein yield even when total expression is high. Of note, the highest expressing C-terminal boundaries for Abl were residues 511 and 512. These residues fall in the regulatory alpha helix *α*|^26^. This helix has been shown to undergo a dramatic conformational change upon binding to the myristoylated N-terminal cap, which introduces a sharp “kink” in residues 516–519. These residues may lead to higher levels of soluble expression by truncating an secondary structural element that is unusually flexible. Indeed, this helix is not resolved in some X-ray structures (PDBID:2E2B)^25^, further suggesting that this helix is less thermodynamically stable than expected. Control replicates of three constructs indicate good repeatability of expression yields in the high-throughput format. This screen suggests that optimization of construct boundaries could potentially further greatly increase yields of poorly expressing kinase domains. Codon optimization for bacterial expression could also increase expression for kinase domains with low yield due to codon bias^49^, as could coexpression with chaperones^50^.

For those kinases that did express, a fluorescence-based thermostability assay indicated that many of the highest-expressing kinases are well folded. An ATP-competitive inhibitor binding fluorescent assay provides qualitative evidence that the 39 kinases that had sufficiently high expression levels to be assayed have a well-formed ATP-binding site capable of binding bosutinib, a small molecule ATP-competitive kinase inhibitor. Taken together, these two experiments demonstrate that our expression protocol produces folded kinases with utility for biophysical experiments and drug design.

The tolerance of these bacterial constructs to many engineered clinical missense mutations suggests a promising route to the high-throughput biophysical characterization of the effect of clinical mutants on anticancer therapeutics. Mutations that did not express well may destabilize the protein, or may increase the specific activity of the kinase. A higher specific activity would require more phosphatase activity, wasting ATP to prevent high levels of phosphorylation that have been hypothesized to cause difficulty expressing kinases without a coexpressed phosphatase in bacteria^21^. Mutations that are destabilizing may show improved expression if coexpressed with more elaborate chaperones such as GroEL and Trigger factor^50^. Mutations that increase the specific activity of the kinase might also express better when combined with an inactivating mutation.

High-throughput automated kinase expression could be combined with enzymatic or biophysical techniques for characterizing the potency of a variety of clinical kinase inhibitors to assess which mutations confer resistance or sensitivity. While the process of engineering, expressing, purifying, and assaying mutants currently takes approximately two weeks, it is possible that new techniques for cell-free bacterial expression^51, 52^ may reduce this time to a matter of days or hours in a manner that might be compatible with clinical time frames to impact therapeutic decision-making.

We hope that other laboratories find these resources useful in their own work.

## Author Contributions

Conceptualization, JDC, DLP, SKA, MI, LRL, SMH, NML, MAS; Methodology, DLP, MI, LRL, SMH, SKA, JDC, NML, MAS; Software, DLP, JDC, SMH; Formal Analysis, SKA, JDC, MI, SMH; Investigation, MI, LRL, SG, CJ, SKA, SMH; Resources, CJ, SG; Data Curation, SKA, MI, LRL, DLP, JMB; Writing-Original Draft, SKA, LRL, DLP, JDC, SG, SMH, MI; Writing - Review and Editing, SKA, JDC, MI, LRL, SHM, SG, CJ, NML, MAS; Visualization, SKA, JDC, MI, SMH; Supervision, JDC, NML, MAS; Project Administration, SKA, JDC, MI, SMH; Funding Acquisition, JDC, SMH

## Acknowledgments

DLP, SMH, LRL, SKA, MI, and JDC acknowledge support from the Sloan Kettering Institute. This work was funded in part by the Marie-Josée and Henry R. Kravis Center for Molecular Oncology, the National Institutes of Health (NIH grant R01 GM121505 and National Cancer Institute Cancer Center Core grant P30 CA008748), the Functional Genomics Institute (FGI) at MSKCC, and a Louis V. Gerstner Young Investigator Award. MAS acknowledges funding support by NIH grant R35 GM119437. The authors are grateful to Gregory Ross (MSKCC) for assistance in preparing the computational infrastructure for selecting clinical point mutants, and to Sarah E. Boyce (current address: Schrödinger, New York, NY) for assistance with multiple stages of this project. We gratefully acknowledge the members of the MSKCC Molecular Diagnostics Service in the Department of Pathology for their efforts in collecting and compiling mutations for Abl and Src kinases used here. We thank the Kuriyan lab for the gift of pCDFDuet1-YOPH plasmid. The authors are grateful to Addgene for their help in making the plasmids generated by this work available to the research community at minimal cost.

## Supplementary Methods

### Large Scale expression and purification protocol for MK14

Large scale expression of MK14 was performed at the QB3 MacroLab (QB3 MacroLab, University of California, Berkeley, CA 94720 [http://qb3.berkeley.edu/macrolab/], a core facility offering automated gene cloning and recombinant protein expression and purification services.

Rosetta2(DE3)pLysS cells (Novagen) were used to co-express MK14 (same plasmid as from the high-throughput kinase expression panel) and 13SA Lamda phosphatase. The cells were grown in 2YT Medium (16 g/L Tryptone, 10 g/L Yeast Extract, 5 g/L NaCl) to OD600 of 0.5 at 37°C. The culture was cooled to 16°C and induced with 0.5 mM IPTG overnight. The cultures were pelleted at 5000 rpm for 30 min and resuspended in 20 mL Nickel buffer A (25 mM HEPES pH 7.5, 10% glycerol, 400mM NaCl, 20 mM imidazole, 5 mM BME) with the following protease inhibitors: 1 *µ*g/mL leupeptin, 1 *µ*g/mL pepstatin, and 0.5 mM PMSF). The resuspended cells were frozen at −80°C.

When ready for purification, the cells were thawed and ruptured using a homogenizer (Avestin C3, 15000psi, 3 passes). The broken cells were pelleted at 15000 rpm for 30 min (SS34 rotor). Clarified lysate was loaded onto a 5 mL HisTrap FF Crude column (GE Healthcare) and washed with Nickel buffer A to remove any unbound material. The protein was eluted with Nickel buffer B (25 mM HEPES pH 7.5, 10% glycerol, 400mM NaCl, 400 mM imidazole, 5 mM BME) and pooled for buffer exchange into Nickel buffer A on a HiPrep 26/10 Desalting Column (GE Healthcare). Rough protein yields were quantified using theorectical extinction coefficients calculated using ProtParam (http://ca.expasy.org/tools/protparam.html). The His tag was cleaved off of MK14 by incubation with TEV protease (25°C, 2 hours, 1:20 mass ratio).

After tag cleavage, the sample was run over a 5 mL HisTrap FF Crude column (GE Healthcare) with Nickel buffer A. The flow-through was collected, concentrated to roughly 5mL using centrifugal concentrators (10 kDA MWCO, Millipore) and loaded onto a HiPrep 16/60 Sephacryl S-200 HR column (GE Healthcare). The sample was equilibrated into Gel Filtration buffer (20 mM Tris-HCl pH 8.0, 150 mM NaCl, 5% glycerol, 1 mM DTT) and fractions containing MK14 were pooled and concentrated (10 kDA MWCO centrifugal concentrators, Millipore). 500 *µ*L aliquots of MK14 were snap frozen in liquid nitrogen and stored at −80°C. Quantification by theoretical extinction coeffcient suggests the final MK14 concentration was roughly 4.0 mg/mL (97 *µ*M), roughly 22.4 mg/L of culture yield.

## Footnotes

1 These targets are, currently: Abl, DDR1, EGFR, HER2, VGFR1/2/3, Alk, Met, BRAF, JAK1/2/3, Btk, Pi3K, CDK4, CDK6, MEK, ROS1, FLt3, IGF1R, Ret, Kit, Axl, TrkB, and mTOR^9^.

2 Parent plasmid is a pET His10 TEV LIC cloning vector and is available on Addgene (Plasmid #78173).

3 Yoph164 phosphatase, engineered to minimize intrinsic affinity for nickel purification resin by the QB3 MacroLab based on parent plasmid pCDFDuet1-YOPH, a gift from the Kuriyan Lab.

